# spectrum_utils: A Python package for mass spectrometry data processing and visualization

**DOI:** 10.1101/725036

**Authors:** Wout Bittremieux

## Abstract

Given the wide diversity in applications of biological mass spectrometry, custom data analyses are often needed to fully interpret the results of an experiment. Such bioinformatics scripts necessarily include similar basic functionality to read mass spectral data from standard file formats, process it, and visualize it. Rather than having to reimplement this functionality, to facilitate this task, `spectrum_utils` is a Python package for mass spectrometry data processing and visualization. Its high-level functionality enables developers to quickly prototype ideas for computational mass spectrometry projects in only a few lines of code. Notably, the data processing functionality is highly optimized for computational efficiency to be able to deal with the large volumes of data that are generated during mass spectrometry experiments. The visualization functionality makes it possible to easily produce publication-quality figures as well as interactive spectrum plots for inclusion on web pages. `spectrum_utils` is available for Python 3.6+, includes extensive online documentation and examples, and can be easily installed using conda. It is freely available as open source under the Apache 2.0 license at https://github.com/bittremieux/spectrum_utils.

## 1 Introduction

Mass spectrometry (MS) is a powerful, high-throughput analytical technique that can be used to identify and quantify molecules in complex biological samples. Because during a typical MS experiment tens of thousands of mass spectra are generated, suitable bioinformatics tools are needed to analyze such large data volumes. MS data processing has traditionally been done using monolithic software tools that aim to provide fully end-to-end solutions from the raw data to the final identification or quantification results. However, because there exist a large variety of experimental set-ups and configurations, such tools necessarily cannot cover all possible use cases.

Instead, customized data analysis workflows are often needed to fully interpret the results of an MS experiment. In recent years several software packages for the general-purpose analysis of MS data in popular scripting languages have been developed. Notable examples include MSnbase [6] for data processing, visualization, and quantification in R; pymzML [1, 10] to efficiently read and process spectra in the mzML format [13] using Python; Pyteomics [7, 12] for a variety of proteomics data processing tasks in Python; and pyOpenMS [20] to expose the rich functionality of OpenMS [19, 22] from C++ to Python.

Here we present the `spectrum_utils` package for MS data processing and visualization in Python. `spectrum_utils` allows the user to easily manipulate mass spectral data and quickly prototype ideas for computational MS projects. A key feature of `spectrum_utils` is its focus on computational efficiency to process large amounts of spectral data. `spectrum_utils` is freely available as open source under the Apache 2.0 license at https://github.com/bittremieux/spectrum_utils.

## 2 Methods & results

The functionality provided by `spectrum_utils` is built around the concept of tandem mass spectrometry (MS/MS) spectra as basic elements. MS/MS spectra can be processed in various ways. Uninformative peaks, such as the precursor ion and its isotopic peaks, and low-intensity noise peaks, can be removed. Further peak filtering can be performed by only retaining the top most intense peaks. Next, peak intensities can be scaled to de-emphasize overly dominant peaks. Possible transformations are root scaling, log scaling, or rank-based scaling. Finally, peaks can be annotated with their peptide fragments, potentially including post-translational modifications (PTMs) at specified amino acid positions, molecules encoded as SMILES strings, or custom annotations (figure 1).

**Figure 1:**
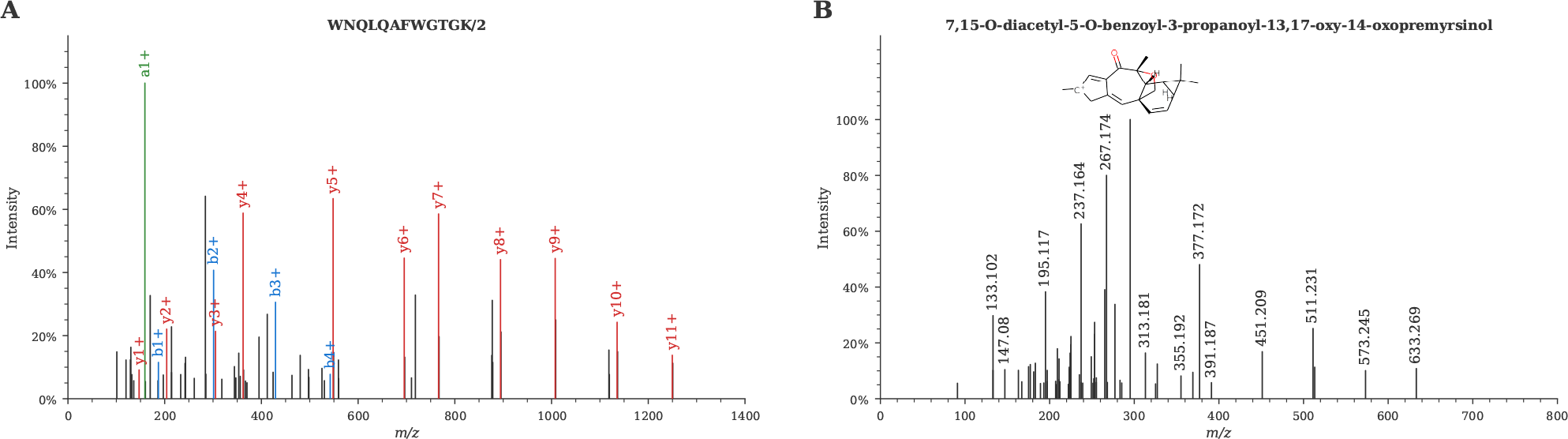
Visualization of (A) a proteomics MS/MS spectrum (mzspec:PXD004732:01650b_BC2-TUM_first_pool_53_01_01-3xHCD-1h-R2:scan:41840) [27] and (B) a metabolomics MS/MS spectrum (mzspec:GNPSLIBRARY:CCMSLIB00000840351) [16]. Fragment peaks can be easily annotated based on a peptide sequence, a SMILES string corresponding to a (sub)structure, or using custom annotations.

An important emphasis is placed on the computational efficiency of the spectrum processing steps. Operations on the peaks of MS/MS spectra are implemented using NumPy [23], a popular Python library for efficient numerical computation. Additionally, Numba [11], a Python just-in-time (JIT) compiler, is used to further speed up many processing steps by compiling Python and NumPy code into efficient machine instructions. Using these techniques for optimized numerical computation, `spectrum_utils` is able to very efficiently process large amounts of MS data (figure 2).

**Figure 2:**
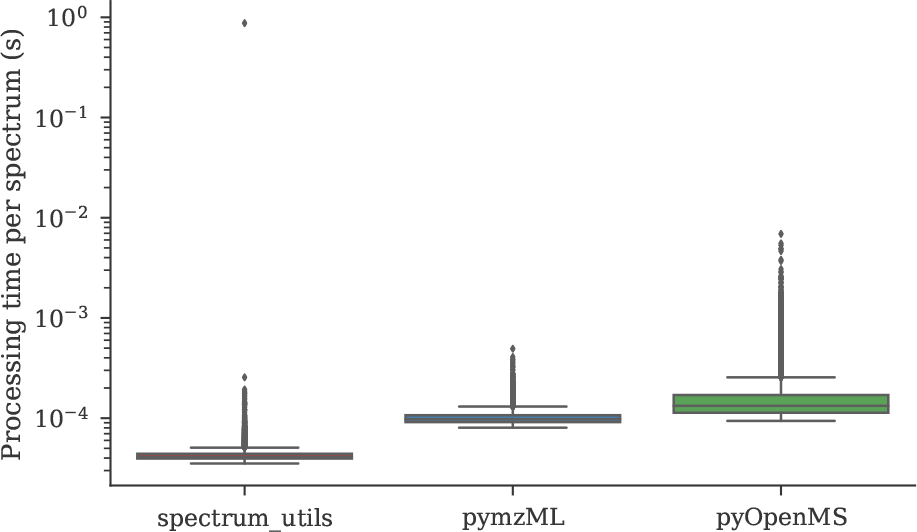
Spectrum processing runtime comparison. The runtime for processing the spectra generated for the iPRG2012 challenge [4] is reported for similar processing steps for `spectrum_utils` (version 0.3.0), pymzML (version 2.4.4) [1, 10], and pyOpenMS (version 2.4.0) [20]. Spectrum processing consisted of fixing the *m*/*z* range, removing precursor ion peaks and low-intensity noise peaks, and scaling the peak intensities by their square root. Note that the significant outlier for `spectrum_utils` is caused by Numba’s JIT compilation [11] of the first method call, allowing subsequent calls to be made very efficiently.

MS/MS spectra can be easily visualized using `spectrum_utils`’s plotting functionality to help in producing publication-quality figures (figure 1). Additionally, a mirror plot can be used to clearly show the similarity between two different spectra, for example, to visualize identification results produced by spectral library searching. The default plotting functionality uses matplotlib [9] to generate high-quality spectrum figures. Alternatively, interactive plotting functionality is available using Altair [24], which is based on the Vega and Vega-Lite grammar of interactive graphics [21]. Interactive plotting is a drop-in replacement for the standard plotting functionality, allowing the user to trivially produce interactive spectrum plots for data exploration or visualization on web pages.

### 2.1 Use cases

We will briefly highlight two use cases to demonstrate the functionality of `spectrum_utils`. First, `spectrum_utils` is used by the ANN-SoLo open modification spectral library search engine [2, 3] to preprocess MS/MS spectra prior to spectral library searching. `spectrum_utils` provides the full functionality needed to preprocess the spectral data, including rounding the peak *m*/*z* values to a common mass resolution, removing the precursor peak and low-intensity noise peaks, and scaling the peak intensities. ANN-SoLo was developed to efficiently perform open modification searches, which typically suffer from a very large search space. In part due to the efficient spectrum processing functionality provided by `spectrum_utils`, ANN-SoLo is able to identify hundreds to thousands of spectra per minute, enabling researchers to perform untargeted PTM profiling via open searches at an unprecedented scale.

Second, the Global Natural Products Social Molecular Networking (GNPS) public data repository and analysis platform uses `spectrum_utils` to produce high-quality spectrum vector graphics. All MS/MS spectra processed by GNPS can be visualized using their universal spectrum identifier [5], as well as the individual spectra in reference spectral libraries compiled using the MSMS-Chooser workflow [25]. Additionally, spectral library identification results obtained using GNPS are visualized using mirror plots.

### 2.2 Code availability

`spectrum_utils` is available for Python 3.6+ and can be easily installed via conda using the Bioconda channel [8]. `spectrum_utils` depends on Numpy [23] and Numba [11] for efficient numerical computation, Pyteomics [7, 12] for peptide fragment ion mass calculations, RDKit [18] for SMILES string handling, matplotlib [9] for static plotting, and Altair [24] and Pandas [15] for interactive plotting.

All code and detailed documentation on how to use `spectrum_utils` is freely available as open source under the Apache 2.0 license at https://github.com/bittremieux/spectrum_utils.

## 3 Conclusion

Here we have presented the `spectrum_utils` package for MS data processing and visualization in Python. Its clearly defined functionality allows `spectrum_utils` to fill an important gap in the Python MS processing ecosystem. For example, `spectrum_utils` does not provide functionality to read spectral data files because there already exist several excellent tools to perform this task. Instead, `spectrum_utils` takes the MS data provided by such tools as input for subsequent processing and visualization. `spectrum_utils` has a well-defined, Pythonic application programming interface, allowing developers to easily harness its powerful functionality in a small number of lines of code. Additionally, it is trivial to switch between the static plotting functionality to produce high-quality spectrum figures to include in scientific manuscripts and the interactive plotting functionality. This interactive plotting functionality can be very powerful during an explorative analysis of MS data or to include dynamic spectrum plots in web resources.

Besides the two use cases highlighted above, `spectrum_utils` has already been used in several other computational MS projects, showcasing its versatile functionality. Among others, it has been used to perform custom analyses of MS data corresponding to an unknown protein sample [17], to prepare MS/MS spectra for processing using deep neural networks [14], and to visualize spectra for scientific publications and presentations. This illustrates how `spectrum_utils` facilitates several MS data processing and visualization tasks and allows developers to quickly prototype ideas and implement diverse computational MS projects.

